# Neural distinctiveness and discriminability in regions of the core network representations support associative unitization

**DOI:** 10.1101/2022.12.20.521200

**Authors:** A. C. Steinkrauss, C. M. Carpenter, M. K. Tarkenton, A.A. Overman, N.A. Dennis

**Affiliations:** The Pennsylvania State University; Elon University

**Keywords:** unitization, associative memory, neural reinstatement, multivariate pattern analyses, neuroimaging

## Abstract

Previous work has suggested unitized pairs behave as a single unit and more critically, are processed neurally different than those of associative memories. The current works examines the neural differences between unitization and non-unitized memory using fMRI and multivoxel analyses. Specifically, we examined the differences across face-occupation pairings as a function of whether the pairing was viewed as a person performing the given job (unitized binding) or a person saying they knew someone who had a particular job (non-unitized binding). The results show that at encoding, the angular gyrus can discriminate between unitized and non-unitized target trials. Additionally, during encoding, the medial temporal lobe (hippocampus and perirhinal cortex), frontal parietal regions (angular gyrus and medial frontal gyrus) and visual regions (middle occipital cortex) exhibit distinct neural patterns to recollected unitized and non-unitized targets. Furthermore, the medial frontal gyrus and middle occipital cortex show greater neural similarity for recollected unitized trials than those of recollected non-unitized trials. We conclude that visually unitized pairs may enhance unitization in older adults due to greater similarity of trials within the same condition during the encoding process.

## 1. Introduction

The ability to form and remember correct associations is essential for recollection in episodic memory. Despite this necessity, the ability to successfully recall associations can be a difficult task in older adults (see Old & Naveh-Benjamin, 2008 for mata-analysis) For example, associating names with faces or stores with addresses can be increasingly difficult as a person ages. The associative deficit hypothesis attributes older adults’ poor associative memory performance to an age-related difficulty in binding together novel pieces of information (Naveh-Benjamin, 2000; Naveh-Benjamin et al., 2004, 2007). Both unitization and visual imagery have been explored as encoding strategies intended to enhance the binding process and thus support older adults’ associative memory (Kondo et al., 2005; Tu & Diana, 2021). The current study seeks to elucidate the neural basis underlying such associative memory strategies in aging individuals. Specifically, we use multivariate pattern classification and neural distinctiveness analyses to investigate whether the advantage afforded by a visual imagery task, used to promote unitization at encoding, supports discriminability and distinctiveness in the neural patterns of activation throughout the associative memory network in older adults.

Unitization is an encoding process used to bind arbitrary items together into a meaningful unit (Ahmad & Hockley, 2014; Bastin et al., 2013; Delhaye et al., 2014; Giovanello et al., 2006; Parks & Yonelinas, 2015; Parra et al., 2009; Quamme et al., 2007). Specifically, the strategy of unitization aims to integrate items in one bound unit which would reduce the involvement of hippocampal-based recollection and increase reliance on perirhinal-based familiarity processing (Ford et al., 2010), implying unitization may behave more similarly to a single item (Graf & Schacter, 1989; J. Haxby et al., 2001; Yonelinas et al., 1999). Most notably, this strategy has examined word-word associations for compound words compared to unrelated word pairs (Ahmad et al., 2015), wherein older adults exhibit enhanced associative recollection for the unitized pairings. Unitization strategies have also been applied to unrelated pairings (Huffer et al., 2022; Memel & Ryan, 2017). For example, asking younger adults to encode face pairs as a “married couple” resulted in better associative memory than two faces encoded as two unrelated individuals (Parks & Yonelinas, 2015). Additionally, Huffer and colleagues (2022), show that unrelated, yet unitized object pairs result in better associative memory compared to a non-unitized presentation of the object pairing in older adults. Importantly, unitization instructions have been shown to improve associative memory in older adults (Kamp et al., 2018; Liu et al., 2022; Overman & Stephens, 2013; Zheng et al., 2016) by increasing both recollection and familiarity-based responding (e.g., Liu et al., 2022).

One often-utilized strategy for inducing unitization has been visual imagery. By visualizing, for example, a color as a feature of an item, it is more likely that a person will create a unitized image of a single object with integrated characteristics (Bastin et al., 2013; Overman & Stephens, 2013; Parks & Yonelinas, 2015; Tu & Diana, 2021; Zheng et al., 2015). For example, Bastin and colleagues (2013) behaviorally, found enhanced associative memory in both older and younger adults using item-color pairings with unitized compared to non-unitized instructions. Using the same paradigm, Zheng and colleagues (2015) found that age differences were reduced in the unitization instruction group due to familiarity-based source recognition. Finally, Tu and Diana (2021) found greater associative memory for younger adults, in all unitized imagery conditions which included a word with color, size, or both color and size. Similarly, face-word pairings that encouraged unitization via visualization and intra-item binding (e.g., imagine a person was a skier) were better remembered than those that encouraged inter-item binding (e.g., the person had simply interacted with a skier) (Overman & Stephens, 2013; Parks & Yonelinas, 2015). It is unknown if older adults will have similar capabilities as young adults to use a unitization-based strategy for an associative memory task, in which they are susceptible to memory declines in. The current work aims to see how in older adults’ visual imagery when using unitization-based strategies can enhance associative memory both behaviorally and neurally. Specifically, this visual imagery will look at face-occupation pairings and imagine those faces as *doing* the job associated with it to then *unitize* the pairing.

Neurally, associative memory has been shown to rely on activity in a common set of regions, including the hippocampus (HC), medial frontal gyrus (MFG), and angular gyrus (AG) (for a meta-analysis see Benoit & Schacter, 2015; Thakral et al., 2017). Specifically, associative memories elicit increases in blood-oxygen-level-dependent (BOLD) activation within the HC for recollection-based associative binding across novel items (Dennis, Bowman, et al., 2014; Dennis, Johnson, et al., 2014; Piekema et al., 2010; Staresina & Davachi, 2010; Yonelinas et al., 2001). Additionally, encoding-based strategies like unitization has been shown to elicit BOLD activation in the perirhinal cortex (PrC) (Haskins et al., 2008; Memel & Ryan, 2017; Staresina & Davachi, 2010), presumably because item-item associations are encoded and retrieved in a manner more similar to that of an item in memory. BOLD activation in the MFG has been found when detecting whether incoming information is congruent with its encoded memory state (Bader et al., 2014). The MFG is also involved in the retrieval of schemas and is engaged when people attempt to establish connections across associative items (Buuren et al., 2014; Kroes & Fernández, 2012; Tse et al., 2011). Additionally, schema retrieval was found during increased BOLD activation in the AG which allows for the encoding and consolidation of new information (Wagner et al., 2015) Wagner and colleagues (2015) found multi-voxel representations of different schema components in the AG during retrieval of novel, but related, information. They concluded that the AG recombines encoded schemas into one integrated memory. In other words, the AG may guide the binding of information from encoding to retrieval by condensing encoded information for better retrieval recollection (Binder et al., 2009; Shimamura, 2011; Wagner et al., 2015). In addition to this core associative network, occipital regions have been consistently activated in visual memory tasks (Dennis et al., 2019; D. C. Park et al., 2004). Specifically, classification accuracy discriminated between visual imagery and stimulus-driven perception in the middle occipital cortex (MOC), showing the MOC is associated to visual imagery (Stokes et al., 2009) and is linked to word-dependent source memory (Park et al., 2013). Park and colleagues (2013) found that for multivariate neural activity in the MOC, contextualized words are processed more like pictures, aiding to the imagery operations of memory processing (Park et al., 2013).

While past work has focused on BOLD differences within this associative network, more recent work is emerging that places an emphasis on understanding memory processing in aging using multivariate analyses. Related to the current work, multivoxel pattern analysis (MVPA) in older adults, has been used to identify unique neural patterns associated with subtle differences in stimulus properties, such as classifying true and false memories (Carpenter et al., 2021; Chamberlain et al., 2022; Dennis et al., 2022), categories like faces and houses (Carp et al., 2011; Johnson et al., 2015) and recollection- and familiarity-based responding vs. responding at chance (Kafkas et al., 2017). Previous work from our own group has found that neural patterns across the encoding network, including those within prefrontal and occipital cortex, can reliably distinguish between different types of associative memory during encoding (Dennis et al., 2019; Elbich et al., 2021). The fact that such subtle differences in neural processing are detected in older adults speaks to the idea that even for similar sets of information, different encoding instructions can result in uniquely represented associations in the encoding network. That is, neural representation for similar information can differ as a function of how information is processed at encoding.

The examination of neural patterns also gives insight on the distinctiveness of different memory processes and how neural representations contribute to and support episodic memory performance (Sommer & Sander, 2022). By examining the distinct neural patterns related to specific memory behaviors, such as associative recollection, we can gain further insight into how unique representations during encoding lead to successful memory. Similar representations within an encoding condition indicate that trials within that condition share overlapping representations of information, which is to be expected when representing several related items from the same category (Kriegeskorte & Kievit, 2013; Sommer & Sander, 2022). However, when information is more distinctly processed in memory, neural patterns will represent this uniqueness, reflected in less overlapping representation within a category of stimuli and instead, more distinct patterns of neural activity. In neuroimaging memory research, different types of associations, such as unitized associations compared to non-unitized associations, rely on different neural processes (LaRocque et al., 2013; Staresina & Davachi, 2010). Knowing how memory is represented and how those representations are related to successful retrieval is important for understanding how we might be able to improve memory processing older adults. It is unknown how older adults’ neural activation may differ in their ability to encode and retrieve information when using a unitization-based strategy versus an associative memory strategy. Specifically using neural distinctiveness analyses in to identify differences in the neural patterns underlying recollection-related activity and underlying unitization for associative memories at each memory stage. Our focus is on regions within the core associative memory network.

The current study aims to examine whether a visual imagery encoding strategy, intended to promote unitization, will result in improved recollection-based associative memory in older adults. The current study also aims to determine whether use of such an encoding strategy enhances the distinctiveness of memory representations during both encoding and retrieval. If the condition promoting unitization is effective in enhancing associative memory in aging, as we have seen in previous work implementing a strategy-based unitization manipulation (Bastin et al., 2013; Delhaye et al., 2014; Diana et al., 2008; Huffer et al., 2022; Quamme et al., 2007), then we expect older adults to have greater recollection in the unitized condition than in the associative memory condition. Neurally, we expect unitized and non-unitized memory for targets to show neural discriminability within the core network, including the HC, PrC, MFG, AG, and MOC. Further, we predict that the unitized encoding strategy pairs would show greater distinctiveness than non-unitized associative memory pairs when taking into account subsequent recollection of the associative information (Kriegeskorte & Kievit, 2013; Simmonite & Polk, 2022; Sommer & Sander, 2022). Our approach was undertaken in order to assess how encoding strategies are processed successfully during an associative memory task.

## 2. Materials and Methods

### 2.1 Subjects

25 older adults were recruited from the Centre County community as a part of a larger study including both younger and older adults. The older adults received monetary compensation for their participation. Participants were screened for history of psychiatric and neurological illness, head injury, stroke, learning disability, medication that affects cognitive and physiological function, and substance abuse. On the day of the study, all participants provided written informed consent for a protocol approved by The Pennsylvania State University Institutional Review Board. All participants were native English speakers or had learned English before the age of 8, with normal or corrected-to-normal vision and were right-handed. All participants had completed high school. All 25 older adults were included in all analyses (*M*_age_= 70.08 years, *SD*_age_= 6.77 years, range = 60-84 years; 14 female, 11 male). Participants identified as white (n = 23), more than one race (n = 1), and preferred not to answer (n = 1), and were all well-educated (*M*_years_ = 16.50, *SD*_years_ = 2.65). All older adults scored above a 27 on the Mini Mental State Examination (MMSE; Folstein et al., 1983; *M* = 29.22, *SD* = 1.02) and below a 3 on the Geriatric Depression Scale (GDS; Yesavage, 1998; *M* = 0.88, *SD* = 1.17).

### 2.2 Regions of Interest (ROIs)

Based upon previous work mentioned above, we restricted our analysis to the angular gyrus (AG), hippocampus (HC), perirhinal cortex (PrC), medial frontal gyrus (MFG), and middle occipital cortex (MOC) (Bader et al., 2014; Binder et al., 2009; Dennis, Bowman, et al., 2014; Dennis, Johnson, et al., 2014; Haskins et al., 2008; Stokes et al., 2009). The ROIs were defined anatomically and created using the human AAL Pickatlas through SPM12 (Folstein et al., 1983) (See Fig. 1).

**Figure 1.**
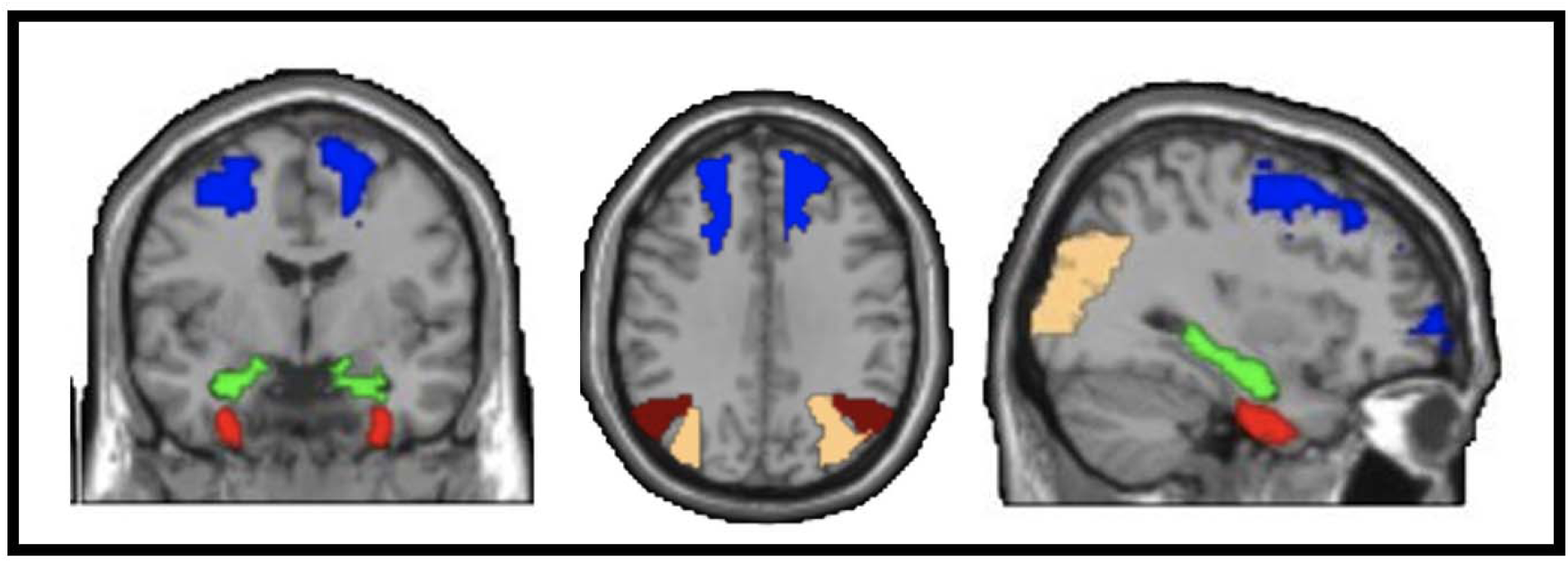
Regions of interest for multivariate classification. Regions defined from AAL pickatlas. Tan = MOC, Dark Red = AG, Blue = MFG, Green = HC, Light Red = PrC. Slice numbers: y = -6; z = 36; x = 32.

### 2.3 Stimuli & Procedure

The current design and stimuli were modified from Overman & Stephens (2013) and previously reported in Ricupero and colleagues (2023) where a young adult sample was analyzed. The experimental stimuli consisted of 144 black and white photographs of faces (see Criss & Shiffrin, 2004 for standardization details) and 144 single-word occupations (ex: pianist; welder), with a majority of the occupations taken from Yovel & Paller (2004) and additional occupations added as needed. During encoding, participants were shown an image of a face and a name tag stating “Hello, I’m [occupation]” (‘doing’ condition) or a speech bubble stating “I know [occupation]” (‘speaking’ condition). Participants were asked to imagine the face-occupation association and remember the pairings using one of two strategies designed to either promote unitization (‘doing’) or simply to create an association (‘speaking’). Specifically, in the ‘doing’ condition participants were asked to imagine the pictured individual performing actions related to the occupation. In the ‘speaking’ condition participants were asked to imagine the pictured individual knowing someone else with the given occupation. During practice, participants were asked to repeat the instructions back and explain what they should be imaging during the two strategy tasks. Participants were prompted by an instruction screen as to which strategy they should employ, with both encoding strategies used in each run. Participants were also asked to indicate, using a four-point button box, how easy or how difficult it was to use the given strategy for each unique face and job pairing (response options included: very difficult, somewhat difficult, somewhat easy, very easy). Finally, the background screen color randomly alternated between either yellow or blue, with each appearing 50% of the time (analyses related to this color manipulation were not included in the current set of analyses).

During encoding trials, participants received a prompt for a duration of 2.5 seconds indicating which of the strategies to use (i.e., speaking). Participants then viewed 9 trials followed by a prompt to use the other strategy (i.e., doing) and then viewed 9 trials. This process repeated itself twice per run. Eleven participants completed this version of task before it was changed to alternate between instructions after 18 trials (N=14), and prompt lasting 5 seconds. There were no repetitions of strategy in each run (i.e., participants saw one doing and one speaking strategy block per run). This change was made in response to feedback from older adults. There were no significant differences in any result between versions. During retrieval trials, participants were presented with both target pairs and recombined lure pairs on a white background. Most face-job pairs were recombined within the same condition^1^. Participants responded during retrieval using a standard Remember-Know-New paradigm (Yonelinas, 2002). They responded ‘Remember’ if they remembered specific details about the face-occupation pair, ‘Know’ if they believed they had seen the pairs together previously, but could not remember specific details of the pair, and ‘New’ if they believed the pair was not presented together previously. Each retrieval run included 36 trials, 24 targets and 12 recombined lures. The order of runs included two encoding runs, followed by two retrieval runs, repeated twice (for a total of four encoding and four retrieval runs). All encoding and retrieval trials were presented for 5 seconds. Participants were provided with practice prior to beginning the task to facilitate learning the two encoding strategies. Participants were given a total of 8 trials at encoding in which they a prompt indicating they should use one of the strategies, then viewed 4 trials, and then were presented with a prompt to use the other strategy and viewed another 4 trials. Following encoding practice, they also had 8 trials of retrieval practice with 6 of those trials being targets. (See Fig. 2).

**Figure 2.**
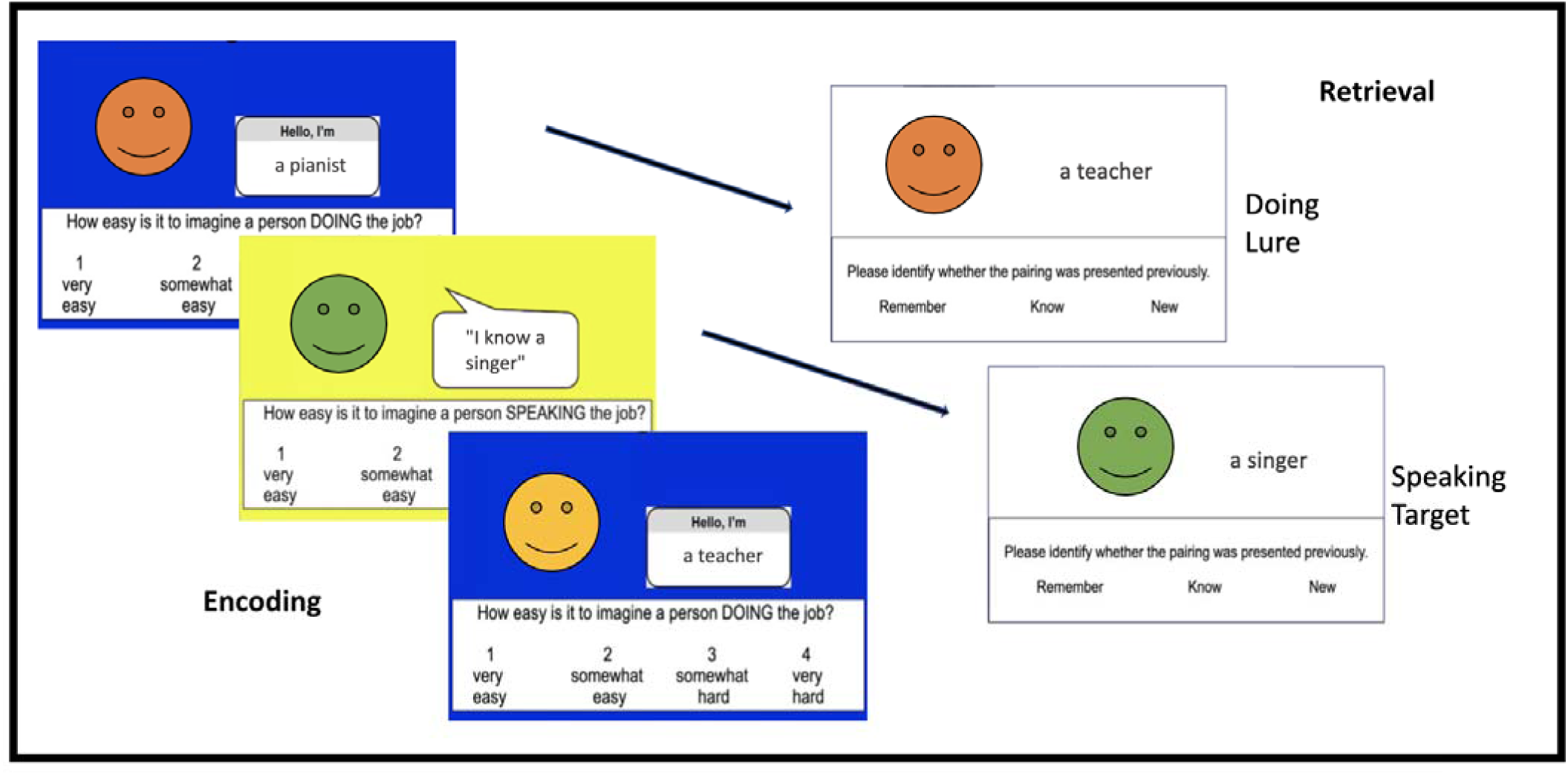
Based on Overman and Stephens 2013, Example stimuli for the unitized and associative memory conditions. Face and job pairings were presented on one of two background colors (yellow or blue). Both the encoding conditions contained blue and yellow trials. At retrieval, the pairings were presented on a white background color. (Note: smile faces are added to replace actual faces for biorxiv, actual images will be reported in the paper)

### 2.5 Image Acquisition

Structural and functional images were acquired using a Siemens 3-T scanner equipped with a 20-channel head coil, parallel to the AC–PC plane. Structural images were acquired with a 2300-msec repetition time, a 2.28-msec echo time, a 256-mm field of view, 192 axial slices, and a 1.0-mm slice thickness for each participant. Echoplanar functional images were acquired using a descending acquisition, a 2500-msec repetition time, a 33-msec echo time, a 192-mm field of view, a 80° flip angle, and 64 axial slices with a 2.0-mm slice thickness resulting in 2.0-mm isotroFpic voxels.

### 2.6 Anatomical Data Processing

A total of 1 T1-weighted (T1w) images were found within the input BIDS dataset. The T1-weighted (T1w) image was corrected for intensity non-uniformity (INU) with N4BiasFieldCorrection (Tustison et al., 2010), distributed with ANTs 2.2.0 (Avants et al., 2008, RRID:SCR_004757), and used as T1w-reference throughout the workflow. The T1w-reference was then skull-stripped with a Nipype implementation of the antsBrainExtraction.sh workflow (from ANTs), using OASIS30ANTs as target template. Brain tissue segmentation of cerebrospinal fluid (CSF), white-matter (WM) and gray-matter (GM) was performed on the brain-extracted T1w using fast (FSL 5.0.9, RRID:SCR_002823, Zhang et al., 2001). Brain surfaces were reconstructed using recon-all (FreeSurfer 6.0.1, RRID:SCR_001847, Dale et al., 1999), and the brain mask estimated previously was refined with a custom variation of the method to reconcile ANTs-derived and FreeSurfer-derived segmentations of the cortical gray-matter of Mindboggle (RRID:SCR_002438, Klein et al., 2017). Volume-based spatial normalization to one standard space (MNI152NLin2009cAsym) was performed through nonlinear registration with antsRegistration (ANTs 2.2.0), using brain-extracted versions of both T1w reference and the T1w template. The following template was selected for spatial normalization: ICBM 152 Nonlinear Asymmetrical template version 2009c [Fonov et al., 2009), RRID:SCR_008796; TemplateFlow ID: MNI152NLin2009cAsym],

### 2.7 Functional data preprocessing

For each of the 8 BOLD runs found per subject (across all tasks and sessions), the following preprocessing was performed. First, a reference volume and its skull-stripped version were generated using a custom methodology of fMRIPrep. Head-motion parameters with respect to the BOLD reference (transformation matrices, and six corresponding rotation and translation parameters) are estimated before any spatiotemporal filtering using mcflirt (FSL 5.0.9, Jenkinson et al., 2002). BOLD runs were slice-time corrected using 3dTshift from AFNI 20160207 (Cox & Hyde, 1997, RRID:SCR_005927). Susceptibility distortion correction (SDC) was omitted. The BOLD reference was then co-registered to the T1w reference using bbregister (FreeSurfer) which implements boundary-based registration (Greve & Fischl, 2009). Co-registration was configured with six degrees of freedom. The BOLD time-series (including slice-timing correction when applied) were resampled onto their original, native space by applying the transforms to correct for head-motion. These resampled BOLD time-series will be referred to as preprocessed BOLD in original space, or just preprocessed BOLD. The BOLD time-series were resampled into standard space, generating a preprocessed BOLD run in MNI152NLin2009cAsym space. First, a reference volume and its skull-stripped version were generated using a custom methodology of fMRIPrep. Several confounding time-series were calculated based on the preprocessed BOLD: framewise displacement (FD), DVARS and three region-wise global signals. FD was computed using two formulations following Power (absolute sum of relative motions, Power et al., 2014)) and Jenkinson (relative root mean square displacement between affines, Jenkinson et al., 2002)). FD and DVARS are calculated for each functional run, both using their implementations in Nipype (following the definitions by Power et al., 2014). The three global signals are extracted within the CSF, the WM, and the whole-brain masks. The head-motion estimates calculated in the correction step were also placed within the corresponding confounds file. The confound time series derived from head motion estimates and global signals were expanded with the inclusion of temporal derivatives and quadratic terms for each (Satterthwaite et al., 2013). Frames that exceeded a threshold of 0.5 mm FD or 1.5 standardized DVARS were annotated as motion outliers. All resamplings can be performed with a single interpolation step by composing all the pertinent transformations (i.e. head-motion transform matrices, susceptibility distortion correction when available, and co-registrations to anatomical and output spaces). Gridded (volumetric) resamplings were performed using antsApplyTransforms (ANTs), configured with Lanczos interpolation to minimize the smoothing effects of other kernels (Lanczos, 1964). Non-gridded (surface) resamplings were performed using mri_vol2surf (FreeSurfer).

### 2.8 Multivariate Pattern Analyses

All analyses were conducted in normalized, MNI space. The anatomical masks were drawn from the AAL pick atlas and were co-registered to a subject’s brain in MNI space. All trials were modeled within each individual trial GLMs. To estimate neural activity associated with individual trials, separate GLMs on unsmoothed data were estimated in SPM12 defining one regressor for each trial at encoding and retrieval (172 total for each phase). An additional 6 nuisance regressors were included in each run corresponding to motion. Whole-brain parameter maps were generated for each trial for encoding and retrieval for each participant. In any given parameter map, the value in each voxel represents the regression coefficient for that trial’s regressor in multiple regression containing all other trials in the run and the motion parameters. These beta parameter maps were concatenated across runs and submitted to CoSMoMVPA toolbox (Oosterhof et al., 2016) for pattern classification (Mumford et al., 2012), and representational similarity/distinctiveness (Haxby et al., 2001) analyses.

#### Neural Pattern Classification

Given our interest in determining how unitization and non-unitized associative memory strategies are discriminated in each region in the associative memory network, classification analyses were conducted to determine if a classifier was able to discriminate between unitization or non-unitized memory target trials in our selected ROIs. Separate classification accuracies were computed between the foregoing trial types at both encoding and retrieval using a support vector machine (SVM) classifier with a linear kernel using all voxels within each ROI (Mumford et al., 2012). Training and testing were performed following an n-fold cross-validation wherein the data was split across the 4 runs. The classifier was trained on 3 runs and tested on the 4th, repeating across different folds. Group-level results were generated from averaging across validation folds from all possible train-data/test-data permutations from the individual participant level. Finally, we tested whether a classifier was significantly able to discriminate neural patterns above chance between the two target types using a one-tailed one-sample t-test for classification accuracy within each ROI. All t-tests were corrected for multiple comparisons using Benjamini-Hochberg corrections and any significance was confirmed with permutation testing (using 10,000 Monte-Carlo simulations). The MVPA was conducted only on target trials, absent of behavior. Classification in the current study begets the idea of pattern distinctiveness compared to chance. The current classification analysis was run to determine how discriminable target trials were from one another. In order to determine how behavior was affected by these trial types, we next conducted pattern distinctiveness analyses accounting for that behavior.

#### Neural Pattern Distinctiveness

Neural distinctiveness metrics examine how distinct neural patterns are from one another in different conditions in order to determine if brain regions are able to discriminate between different conditions or stimuli (Haxby et al., 2001; Kriegeskorte, 2008; LaRocque et al., 2013). Pattern distinctiveness analyses were conducted in the current study to examine the representation of stimuli associated with unitization and non-unitized memory on subsequently recollected targets. Thus, trials will be unequal between conditions and participants. Recollected targets are encoded trials that are subsequently responded to with the subjective response of ‘Remember’ during retrieval, indicating a detailed memory for that pair. For the current analyses, the similarity of activation patterns across different trials of the same category was calculated (within-category correlation, a measure of reliability; Simmonite & Polk, 2022) in each participant for participants subsequently recollected trials (e.g., correlation between all beta parameter maps for unitized subsequently recollected trials correlated individually to all beta parameter maps for all other unitized subsequently recollected trials at encoding) and a between-category similarity score for each participant’s subsequently recollected trials (e.g., one beta parameter map for a unitized subsequently recollected trial correlated individually to all beta parameter maps for non-unitized subsequently recollected trials, done for all unitized trials individually and then averaged) (Haxby et al., 2001). As described in Simmonite and Polk (2022), increased within-category similarity indicates greater reliability and consistency of neural patterns within a given condition. Next, an overall distinctiveness score was calculated by taking the mean of the unitized-within and non-unitized-within similarities scores and subtracting the between similarity score. This value was then compared to 0 using one-sample t-tests, for each ROI, to determine if distinctiveness differed from chance. To further examine whether the discrete distinctiveness was driven by within-category differences, the within-category similarity for the unitized and non-unitized conditions were compared using paired t-tests within each ROI. This process was repeated for both encoding and retrieval trials separately. (See Fig. 3 for an example).

**Figure 3.**
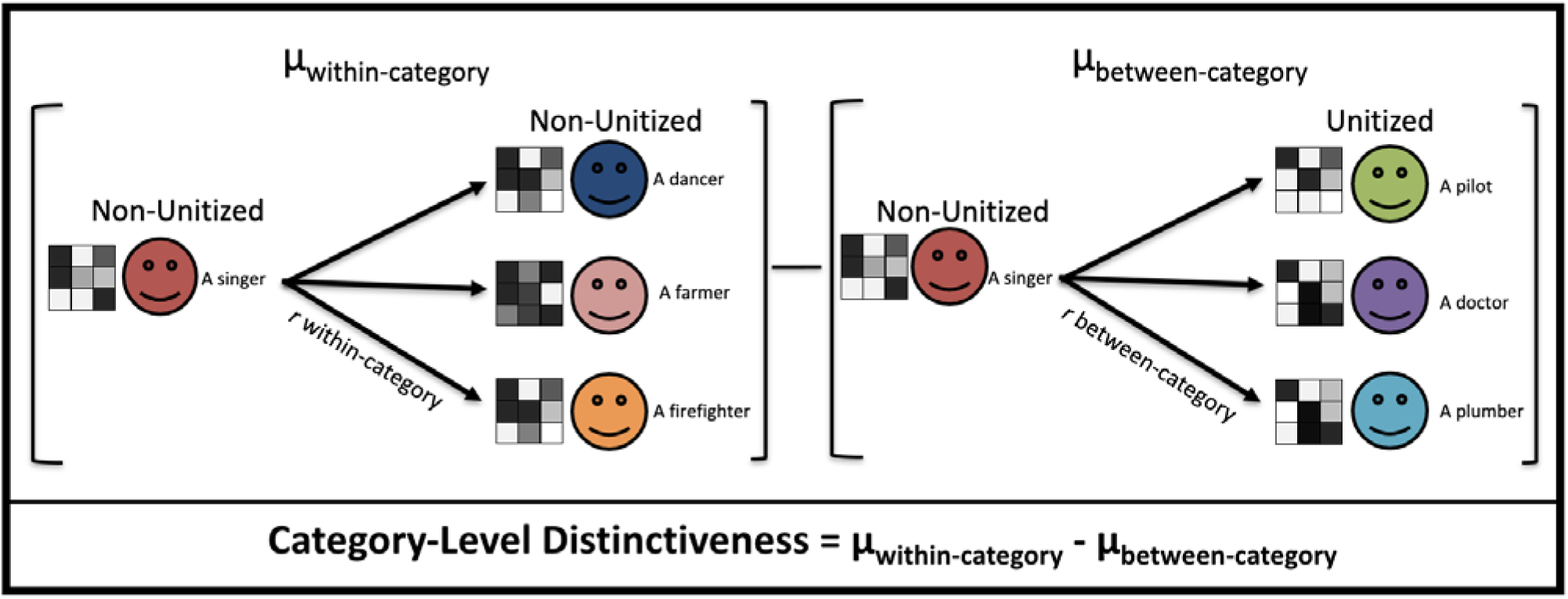
Based on Haxby et al. 2001, calculation of neural distinctiveness. Calculations were repeated twice; once for encoding trials and once for retrieval trials. Distinctiveness was calculated by taking the within-condition similarity correlations for all trials within a condition subtracted by the between-condition similarity correlation for all trials between the condition. (Note: smile faces are added to replace actual faces for biorxiv, actual images will be reported in the paper)

## 3. Results

### Behavioral Results

Overall hit rates across conditions were relatively high, (unitized M= 0.79 SD=0.11 and non-unitized M=0.68, SD=0.15), as were correct rejection rates (unitized M=0.54, SD=0.21 and non-unitized M=0.52, SD=0.16). A series of ANOVAs were run between the unitized and non-unitized memory trials for recollected hits, familiarity hits, and correct rejections (CR), with version of the task included as a covariate. With respect to recollected hit rates, there was a main effect of condition, *F*(1,23) = 61.45, *p* < .001, *d* = .728, such that the unitized memory trials had greater recollected hits (M=0.54 SD=0.20) compared to the non-unitized memory trials (M=0.41 SD=0.17). The main effect of version, *F*(1,23) = 0.41, *p* = .53, *d* = .018, and the interaction between version and condition, *F*(1,23) = 0.03, *p* = .88, *d* = .001, were not significant. With respect to familiarity hits, we found no main effects between unitized (M=0.25 SD=0.16) and non-unitized (M=0.27 SD=0.13) memory trials with version as a covariate, condition, *F*(1,23) = 0.76, *p* = .39, *d* = .032, version, *F*(1,23) = 0.12, *p* = .73, *d* = .005, or a condition by version interaction, *F*(1,23) = 0.36, *p* = .55, *d* = .015. Similarly, for CR, we found no main effects for condition, *F*(1,23) = 0.13, *p* = .72, *d* = .006, version, *F*(1,23) = 2.21, *p* = .15, *d* = .088, or a condition by version interaction, *F*(1,23) = 0.04, *p* = .85, *d* = .002.

### Classification Results

To examine whether classifiers were able to significantly discriminate between our two target conditions, two multivoxel pattern analyses were run. The first classification analysis attempted to classify all unitized and non-unitized memory trial targets at encoding, and the second to classify all unitized and non-unitized memory trial targets at retrieval. Comparing classification of unitized and non-unitized memory trial targets at encoding against chance (50%) showed that in the AG, the classifier accuracy was significantly above chance (AG: *M_accuracy_*= 0.540, *t*(24) = 1.94, *p* < .05). To confirm significance, permutation testing was conducted, using 10,000 Monte-Carlo simulations performed within the AG. In doing so, the AG maintained significance, *p =* .02. Furthermore, in addition to identifying regions showing signals for unitized targets that were discriminable from those of non-unitized targets, we also tested whether there were univariate activation differences in the AG. In the AG, the region showing above-chance classification performance, we computed an ANOCVA with classification accuracy, including mean univariate activation of unitized targets versus baseline and non-unitized targets versus baseline as nuisance covariates. The two covariates did not show a significant difference for target classification (*p*’s > .7). No other ROIs were significant at encoding (HC: t(24)= -0.97, p = .83; MFG: t(24)= 0.544, p = .30; MidOcc: t(24)= 1.304, p = .10; PrC: t(24)= 0.153, p = .44. At retrieval, no regions of interest showed significant classification of unitized and non-unitized memory against chance (AG: t(24)= 1.57, p = .06; HC: t(24)= -0.52, p = .70; MFG: t(24)= 1.53, p = .07; MidOcc: t(24)= 0.57, p = .29; PrC: t(24)= 0.47, p = .32). See Table 1 for all means and standard deviations.

**Table 1.** Means and standard deviations of each ROI for classification analyses during encoding and retrieval.

| ROI | Encoding |  | Retrieval |  |
| --- | --- | --- | --- | --- |
|  | Mean | Standard Deviation | Mean | Standard Deviation |
| AG | 0.540 | 0.104 | 0.517 | 0.053 |
| HC | 0.479 | 0.109 | 0.493 | 0.064 |
| MFG | 0.509 | 0.084 | 0.513 | 0.044 |
| MOC | 0.526 | 0.098 | 0.508 | 0.069 |
| PrC | 0.502 | 0.072 | 0.505 | 0.058 |

### Encoding Distinctiveness

In order to examine neural discriminability related to successful memory at encoding, a neural distinctiveness calculation (Haxby et al., 2001) was conducted for unitized and non-unitized memory subsequently recollected targets (within-category similarity minus between-category similarity) for each encoding condition. At encoding, overall distinctiveness of recollected targets, collapsed across condition, was significantly greater than 0 within all ROIs [AG: *t*(24) = 4.43, *p <* .001; HC: *t*(24) = 2.98, *p <* .01; MFG: *t*(24) = 5.49, *p <* .001; MOC: *t*(24) = 4.79, *p* < .001; PrC: *t*(24) = 2.31, *p <* .05]. The within-condition similarity scores associated with both conditions were then compared to determine if one condition showed higher similarity in a given region compared to the other condition. A direct comparison of within-condition similarity between conditions found significant differences in the MFG and MOC [MFG: *t*(24) = 2.72, *p <* .05, 95% CI[0.01, 0.04]; MOC: *t*(24) = 2.11, *p <* .05, 95% CI[0.00, 0.09]] such that the unitized condition exhibited greater within-condition similarity than the non-unitized condition in the MFG (*M*_unit_ = 0.09 *SD*_unit_ = 0.05, *M*_assoc_ = 0.07 *SD*_assoc_ = 0.04) and MOC (*M*_unit_ = 0.31 *SD*_unit_ = 0.16, *M*_assoc_ = 0.27 *SD*_assoc_ = 0.12). There were no other significant differences in any other ROIs [HC: t(24)=0.97, p=.34, 95% CI[-0.02, 0.06]; PrC: t(24)=0.53, p=.60, 95% CI[-0.02, 0.04]; AG: t(24)=0.13, p=.90, 95% CI[-0.03, 0.04]] (See Fig. 4).

**Figure 4.**
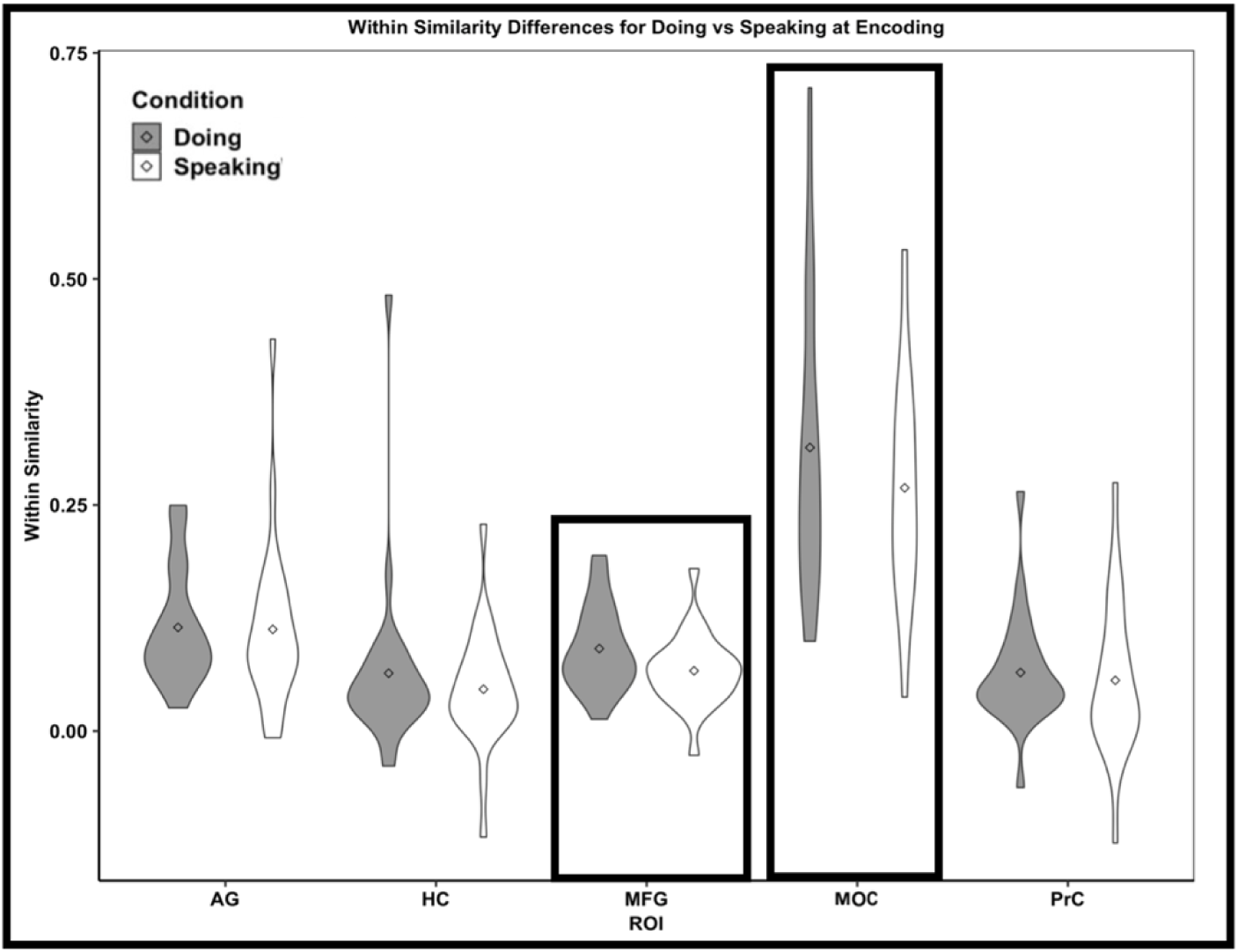
Within-similarity between the two encoding conditions in the AG, HC, MFG, MOC, and PrC. Neural within-similarity scores significantly different between doing versus speaking conditions indicated in a black box; *p’*s < 0.05.

### Retrieval Distinctiveness

In order to examine neural discriminability between two successful retrieval conditions, a neural distinctiveness calculation was conducted for unitized and non-unitized memory recollected targets (within category similarity minus between category similarity). At retrieval, overall distinctiveness of subsequently recollected targets, collapsed across condition, was not significantly greater than 0 within any ROIs [AG: t(24)= -1.17, p=.87; HC: t(24)= .62, p = .27; MFG: t(24)= -.02, p = .51; MOC: t(24)= -1.00, p = .84; PrC: t(24)= 1.06, p = .15]. Since no region of interest was significantly distinct above 0, these conditions were not compared to determine if one condition showed higher similarity in a given region compared to the other condition.

### Exploratory Analysis

As an exploratory analysis, correlation coefficients between distinctiveness in all ROIs were computed in order to examine cross-region relationships in distinctiveness measures during encoding (see Supplemental Fig. 1 for correlations and scatterplot) (see Simmonite & Polk, 2022 for similar analyses). Across all participants, there was a significant positive relationship between the AG and MFG [*r*(23) = .64, *p* < .001.], AG and PrC [*r*(23) = .71, *p* < .001.], MOC and HC [*r*(23) = .48, *p* < .05.], HC and PrC [*r*(23) = .40, *p* < .05.], and MFG and MOC [*r*(23) = .47, *p* < .05.]. The relationship between all other ROIs were not significant (all *p*’s > .05: AG and HC [*r*(23) = .24, *p* = .26], AG and MOC [*r*(23) = .32, *p* = .12], HC and MFG [*r*(23) = .13, *p* = .54], MFG and PrC [*r*(23) = .31, *p* = .14], and MOC and PrC [*r*(23) = .28, *p* = .17.]). All significant findings were confirmed using Benjamini-Hochberg (BH) corrections.

## 4. Discussion

The goal of the current study was to examine the underlying neural mechanisms of unitization, induced through the manipulation of encoding instructions that employed visual imagery. During encoding, participants were given two strategies, one of which promoted unitization of face-occupation pairs through intra-item binding and visualization, while the other, non-unitized encoding strategy, used visualization to promote inter-item binding. Specifically, in the unitized encoding strategy, participants saw a face-occupation pairing and were prompted to imagine the person they were viewing as doing the occupation. In contrast, in the non-unitized encoding strategy participants saw a face-occupation pairing and were prompted to imagine the person they were viewing as knowing someone with the occupation. In alignment with previous work using this paradigm with younger adults (Ricupero et al., 2023), and older adults (Overman & Stephens, 2013), as well as similar studies in older adults, (e.g., Bastin et al., 2013; Delhaye et al., 2014; Diana et al., 2008; Parks & Yonelinas, 2015; Quamme et al., 2007) we found that the unitized condition resulted in greater recollected hits than the non-unitized condition in our sample of older adults. Results support the idea that intra-item binding leads to enhanced associative memory and highlights the ability of older adults to take advantage of such a strategy.

We hypothesized that across both memory phases, information processed within the unitized encoding condition would show greater neural discriminability and neural distinctiveness than information in the non-unitized encoding condition (Kriegeskorte & Kievit, 2013; Simmonite & Polk, 2022; Sommer & Sander, 2022). However, only within the AG at encoding, did MVPA exhibit greater than chance classification between the two encoding conditions. This is consistent with prior work in which AG activation is associated with the presence of schemas during encoding and posited to aid participants to consolidate new information (Binder et al., 2009; Shimamura, 2011; Wagner et al., 2015), including contributing to the encoding of a schema into a bound memory (Wagner et al., 2015). In the current study, classification within the AG may have reflected the difference in processes used to encode an integrated schema of person and their occupation as opposed to the mere association amongst a face and an occupation of a separate individual. This differs from previous aging work in our lab (Dennis et al., 2019) in which it was found that neural patterns across the encoding network can reliably distinguish between two different types of visual associations. One difference across results may be that the current study used similar configurations for both conditions even though the visual associations created by the encoding instructions differed. Specifically, our prior study found that when encoding face-scene associations, regions within prefrontal and occipital cortex, as well as the perirhinal cortex, reliably discriminated between item-item and item-context presentations in older adults. While both the current and prior study controlled for the content of information across associative encoding conditions, our previous study included a greater configural difference across conditions, whereas a similar visual layout was used across encoding conditions in the current study (see Fig. 1). Taken together, the results suggest that encoding processing differences alone may not be enough to promote distinct neural patterns during associative memory in aging within medial temporal regions including the hippocampus and perirhinal cortex. Rather, it may be that in order to represent information uniquely across different encoding conditions, there must be both configural differences and differences in visual encoding between conditions. This aligns with computational modeling work that has suggested a primary age difference in associative memory may be a decrease in the uniqueness of encoded features that enable accurate discrimination of old versus new information (Stephens & Overman, 2018). Further work should continue to explore how encoding strategy and configural differences across encoding conditions supports such neural discriminability.

Similar to the concept of neural discriminability across encoding conditions, the investigation of neural distinctiveness, related to behavioral outcomes (Haxby et al., 2001), allows us to investigate whether neural patterns of successfully recollected unitized and non-unitized memories are presented uniquely both within and across study conditions. In the current study, subsequently recollected targets from both conditions exhibited above-chance distinctiveness across all regions of interest (HC, PrC, MFG, AG, and MOC), indicating that neural patterns associated with these associative pairs are distinct from one another within each condition, during encoding. The fact that recollected trials were only distinct from one another at encoding, and not retrieval, suggests that older adults utilize encoding based strategies and schemas to later benefit their recollection for face-occupation pairings. An exploratory analysis examining the correspondence of distinctiveness across regions found significant cross-region relationships in distinctiveness for the two encoding conditions. Specifically, the HC and PrC had a positive relationship between each other’s distinctiveness indicating these two MTL regions elicit similar activation during encoding across conditions for recollected target trials. Additionally, the AG and MFG, AG and PrC, MOC and HC, and MFG and MOC were also positively correlated to each other with regard to distinctiveness amongst recollected target trials during encoding. Together the results indicate that, not only do MTL regions exhibit similar distinct activation between one another, but so do prefrontal and visual regions as well. This finding suggests that for older adults, processing within different regions in the brain are related to each other when discriminating between recollected trials.

When directly comparing pattern similarity of the unitized condition to that of the non-unitized condition, the reliability of neural activity of each condition was significantly different from each other in the MFG and the MOC, such that the unitized condition exhibited greater reliability (greater within-condition similarity of trials) than the non-unitized condition. This suggests that when processing targets that were later correctly recollected, participants encoded the trials across the unitized condition in a more similar manner than they did those within the non-unitized condition. Since analyses were completed on targets that were later recollected, greater reliability related to within-category activation may likely underscore less detailed-based memory traces. Additionally, this suggests that the MFG and MOC may play a role in distinguishing between associations that are encoded in a different manner. The MFG has been shown to aid in the detection of congruent information from encoding to retrieval as well as to aid in supporting the retrieval of schemas (Kroes & Fernández, 2012; Tse et al., 2011). Since neural patterns in the MFG associated with the unitized condition were more similar to other recollected trials in the same condition, it further supports the idea that the MFG contributes to the encoding of integrated schemas, such as imagining a relationship between a person and their occupation. Since the unitization condition in the current study asked older adults to use schemas to bind the face-occupation pairing, the additional detail associated with the schemas may have led to greater recollection in the unitized condition compared to the non-unitized condition. However, further work such as a questionnaire to identify strategy usage may be necessary to determine how these schemas affected memory performance.

In the unitized condition, it was emphasized that they should be imagining and visualizing the face-occupation pairing especially to a greater extent than the non-unitized condition. Greater unitized trial activation in the MOC supports the idea that older adults engaged in the imagery strategy that they were instructed to use, since the MOC is crucial for visual imagery (e.g., Pidgeon et al., 2016; Stokes et al., 2009). In this study, unitization was designed to be implemented as a top-down mechanism, aiding in the binding of two unrelated items (face-occupation pairing) into a single representation using stored schemas associated with occupation actions. Invoking occupation schemas, combined with generating mental imagery of the actions associated with the occupation, allowed participants to draw on pre-existing information during associative encoding, in a manner similar to that used in previous work (Bastin et al., 2013; Memel & Ryan, 2017). Bastin and colleagues (2013) asked participants to form visual associations and found that older adults had an increased associative memory compared to the non-unitized strategy. In a similar manner, Memel and Ryan (2017) used a visual integration task to bind objects and contexts, in which they placed objects within a logical scene in order to promote schema-related unitization during encoding. This presentation resulted in better associative recollection and increased activation in the prefrontal cortex within their older participants. They concluded visual integrated unitization may be mediated by recollective based encoding processes for older adults (Memel & Ryan, 2017).

## 5. Limitations/Future Directions

While the current findings extend our knowledge of the role of schemas during encoding for older adults, there were some limitations to the study. Specifically, we cannot be certain that the unitized condition is truly being bound as a single unit, only that behavioral differences across encoding conditions suggest stronger memory for the unitization-based strategy. The unitized condition can only be concluded as being different from the non-unitized associative condition, in a manner that is beneficial to older adults’ memory performance. Future work should aim to identify whether these differences in encoding strategies are due to unitization of unrelated information into a singular bound unit. Additionally, further research is needed to determine if a visualization unitization strategy is the most effective strategy to achieve unitized binding. In addition, the benefit of unitization has often been associated with greater use of familiarity, particularly in older adults (Delhaye & Bastin, 2018; Diana, Ranganath & Yonelinas, 2008). However, recent work in the realm of neuroimaging has also indicated that recollection may also be important to mediating the effect of unitization on memory (Memel & Ryan, 2017). In the current task, familiarity-based hit counts were very low and thus we chose to focus solely on recollection-based hits. Future work should aim to investigate and disentangle the role of familiarity and recollection in unitization in terms of neural distinctiveness.

## 6. Conclusions

Overall, the results show that older adults are able to benefit from associative encoding instructions that emphasize unitized binding and visualization, exhibiting greater recollection of targets in the unitized compared to non-unitized condition. Contrary to our predictions regarding neural discriminability, only the AG was able to discriminate between unitized and non-unitized targets during encoding, suggesting that more contextual differences may be required for older adults to recruit other associative memory regions (Dennis et al., 2019). In contrast, we found that neural patterns related to unitized and non-unitized associations were distinct from one another within the core associative network and visual imagery regions. Specifically, when comparing the similarity of unitized versus non-unitized trials, the MOC and MFG exhibited greater reliability at encoding within neural representations of later recollected unitized pairs compared to non-unitized pairs. Overall, the results suggest that unitization may help older adults bind new information together into single units when using visual imagery strategies during encoding, which may be in part due to the ability of visual imagery regions to distinguish between types of associative information.

## Acknowledgements

We would like to thank Courtney Gerver, Harini Babu, Kayla McGraw, Valerie Goodwin, Chloe Hultman and Min Sung Seo for helping with data collection and analysis. This paper was supported by the National Institutes of Health under grant R15 AG052903 awarded to A.A.O and N.A.D.

## Availability of Data and Materials

The datasets generated and analyzed during the current study are available from the corresponding author on reasonable request. None of the experiments were preregistered.

## Footnotes

1 Due to an error in programming, a subset of lures were recombined between (rather than within) condition. All between condition lures were removed from behavioral and imaging analyses.

## Supplemental Figure

**Supplemental Figure 1.**
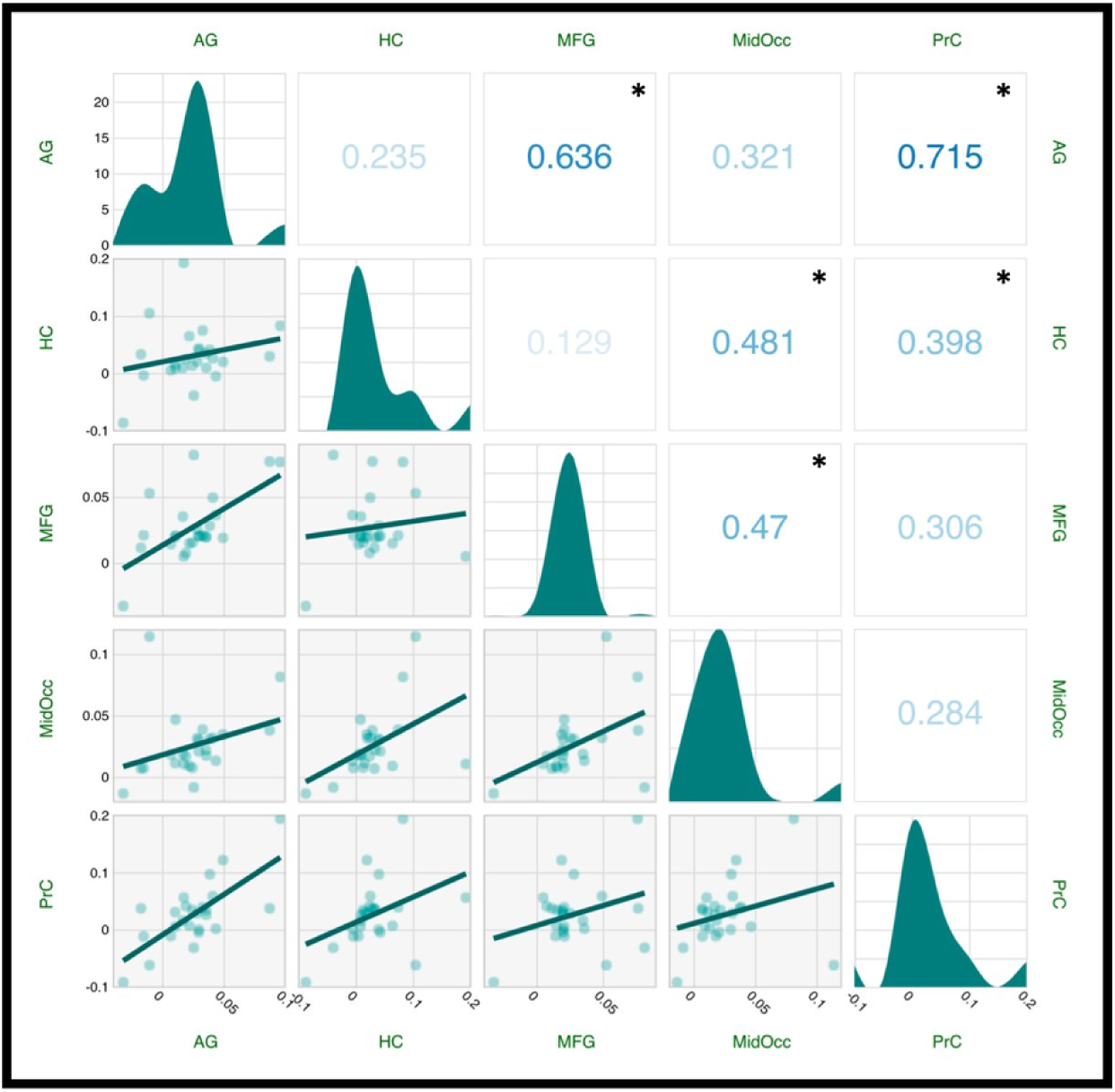
Correlations of neural distinctiveness scores between each AG, HC, MFG, MOC, and PrC during encoding. * indicates regions significantly correlated to each other (*p’*s < .05)

