## Supplemental Figure 1 for "Neural distinctiveness and discriminability in regions of the core network representations support associative unitization"

**Figure legends**

Supplemental Figure

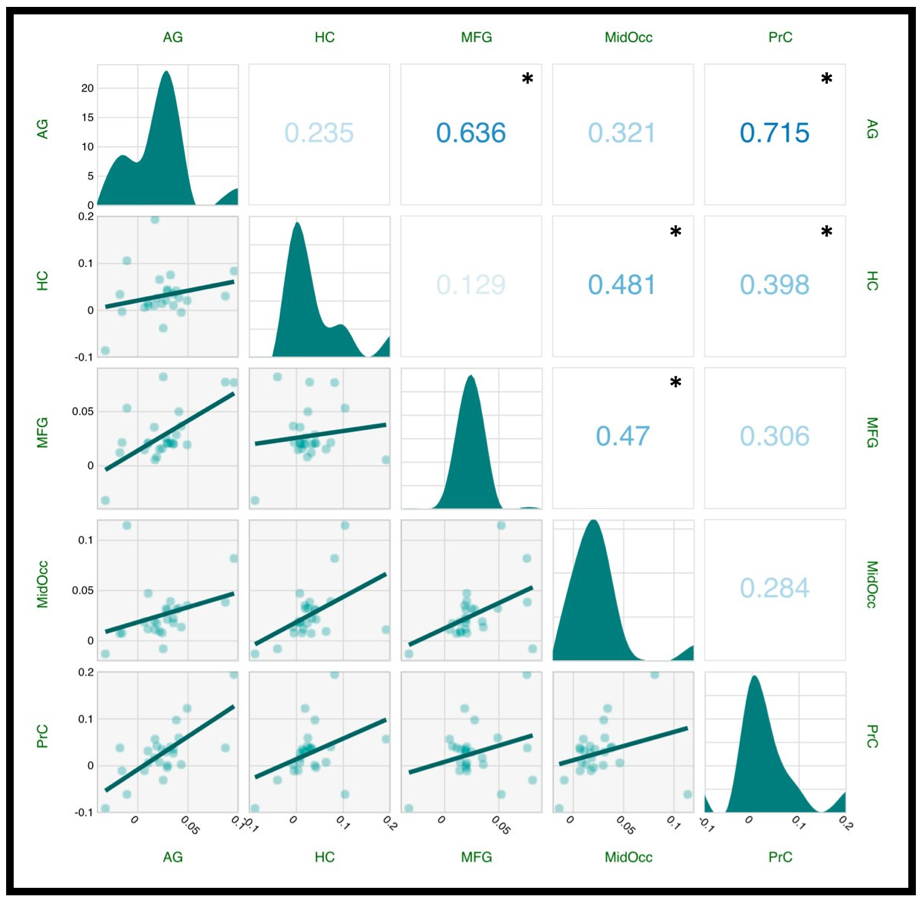

Supplemental Figure 1. Correlations of neural distinctiveness scores between each AG, HC, MFG, MOC, and PrC during encoding. * indicates regions significantly correlated to each other (*p'*s < .05)
